# Multi-parametric characterization of brain-wide hemodynamic and calcium responses to sensory stimulation in mice

**DOI:** 10.1101/2021.11.08.467725

**Authors:** Zhenyue Chen, Quanyu Zhou, Xosé Luís Deán-Ben, Irmak Gezginer, Ruiqing Ni, Michael Reiss, Shy Shoham, Daniel Razansky

## Abstract

Modern optical neuroimaging approaches are expanding our ability to elucidate complex brain function. Diverse imaging contrasts enable direct observation of neural activity with functional sensors along with the induced hemodynamic responses. To date, decoupling the complex interplay of neurovascular coupling and dynamical physiological states has remained challenging when employing single-modality functional neuroimaging tools. We devised a hybrid fluorescence optoacoustic tomography (FLOT) platform combined with a custom data processing pipeline based on statistical parametric mapping, accomplishing the first simultaneous noninvasive observation of both direct and indirect brain-wide activation patterns with optical contrast. Correlated changes in the oxy- and deoxygenated hemoglobin, total hemoglobin, oxygen saturation and rapid GCaMP6f fluorescence signals were observed in response to peripheral sensory stimulation. While the concurrent epifluorescence served to corroborate and complement the functional optoacoustic observations, the latter further aided in decoupling the rapid calcium responses from the slowly varying background in the fluorescence recordings mediated by hemodynamic changes. The hybrid imaging platform expands the capabilities of conventional neuroimaging methods to provide more comprehensive functional readings for studying neurovascular and neurometabolic coupling mechanisms and related diseases.

## Introduction

Functional neuroimaging has become a primary tool in neuroscience and research into neurodegenerative diseases. Blood-oxygen-level-dependent (BOLD) functional magnetic resonance imaging (fMRI) is the mainstay for noninvasive mapping of brain function^1^. However, interpretation of activity-related BOLD signals is challenged by their dependence on multiple factors such as blood flow, blood volume, metabolic rate of oxygen and baseline physiological state^2^. Functional ultrasound imaging can provide additional information on the blood flow^3^. However, the blood flow changes similarly represent an indirect correlate of the underlying neural dynamics, which is insufficient for fully characterizing the complex interplay between neural activity and its accompanying hemodynamics.

Optical imaging approaches arguably provide the most powerful means for retrieving functional and molecular information underlying brain activity. The emergence of GCaMP-type and other genetically encoded calcium indicators has facilitated fluorescence-mediated sensing of neural activation^4^, and has for example been widely employed to characterize resting state activity and stimulus-evoked responses in the mouse cortex^5^. In addition, hemoglobin exhibits distinct absorption spectra in its oxygenated (HbO) and deoxygenated (HbR) states^6^, which can be visualized with functional optoacoustic (fOA) tomography across 3D space and time, enabling the mapping of multiple hemodynamic parameters across the entire murine brain, not accessible with other modalities^7–9^. Imaging modalities based on optical contrasts can thus potentially massively advance our knowledge on the mechanisms of neurovascular coupling, that is, if the mapping of both the neuronal activity directly as well as its associated hemodynamic changes could be realized simultaneously. However, a detailed characterization and rigorous side-by-side validation of hemodynamic fOA readings against well-established, *direct* optical neuroimaging methods is currently lacking.

Here, we introduce a hybrid fluorescence and optoacoustic tomography (FLOT) platform for concurrent multi-parametric characterization of brain-wide hemodynamic and calcium responses to sensory stimulation in mice. A custom data processing pipeline inspired by statistical parametric mapping (SPM) was further developed to facilitate the functional data analysis. Simultaneous imaging of calcium and hemodynamic responses to electrical paw stimulation was performed in GCaMP6f mice, for which the GCaMP signals were shown to reflect changes in intracellular calcium corresponding to neural spiking activity^5,10^. To the best of our knowledge, this represents the first successful observation of activation patterns (direct and indirect responses simultaneously) in the entire mouse cortex with optical contrast. The use of near-infrared (NIR) wavelengths in fOA imaging further enabled to reach deeper brain regions thus averting cross-talk between calcium and hemodynamic signals.

## Results

### The hybrid fluorescence and optoacoustic tomography (FLOT) system

The dedicated hybrid FLOT imaging system employs a flexible fiberscope for capturing fluorescence images in epi-illumination mode with ^~^44 μm pixel resolution. The distal end of the fiberscope was inserted into the central cavity of a spherical matric array transducer used for acquiring three dimensional (3D) fOA data with nearly isotropic spatial resolution which was estimated to be 113 μm using microsphere measurements, slightly degrading at the periphery FOV (Fig. 1**a** and Fig. S1)^11,12^. Both modalities provided a 12 × 12 mm field of view (FOV) along the lateral x-y plane covering the entire mouse cortex. Trains of unipolar rectangular electrical pulses with 5 ms duration and 1.0 mA intensity were applied to the left hindpaw of Thy1-GCaMP6f mice. The trains consisted of 32 pulses with stimulus frequency of 4 Hz (i.e., 8 s total duration) repeated every 82 s (Fig. 1**b**). Five wavelengths (700, 730, 755, 800 and 850 nm) were optimally selected for fOA to avoid significant spectral coloring while ensuring a small condition number of the spectral unmixing matrix (Fig. 1**c**). The unmixed hemodynamic components of HbO and HbR are shown in Fig. 1**d**. Subsequently, both fluorescence and fOA data were analyzed with a dedicated data processing pipeline (Fig. 1**e**). We employed regressors for analyzing the hemodynamic and calcium responses by convolving the stimulation pulse train with a hemodynamic response function (HRF) and a GCaMP calcium response function (CRF), respectively (see Methods for a detailed description).

**Figure 1.**
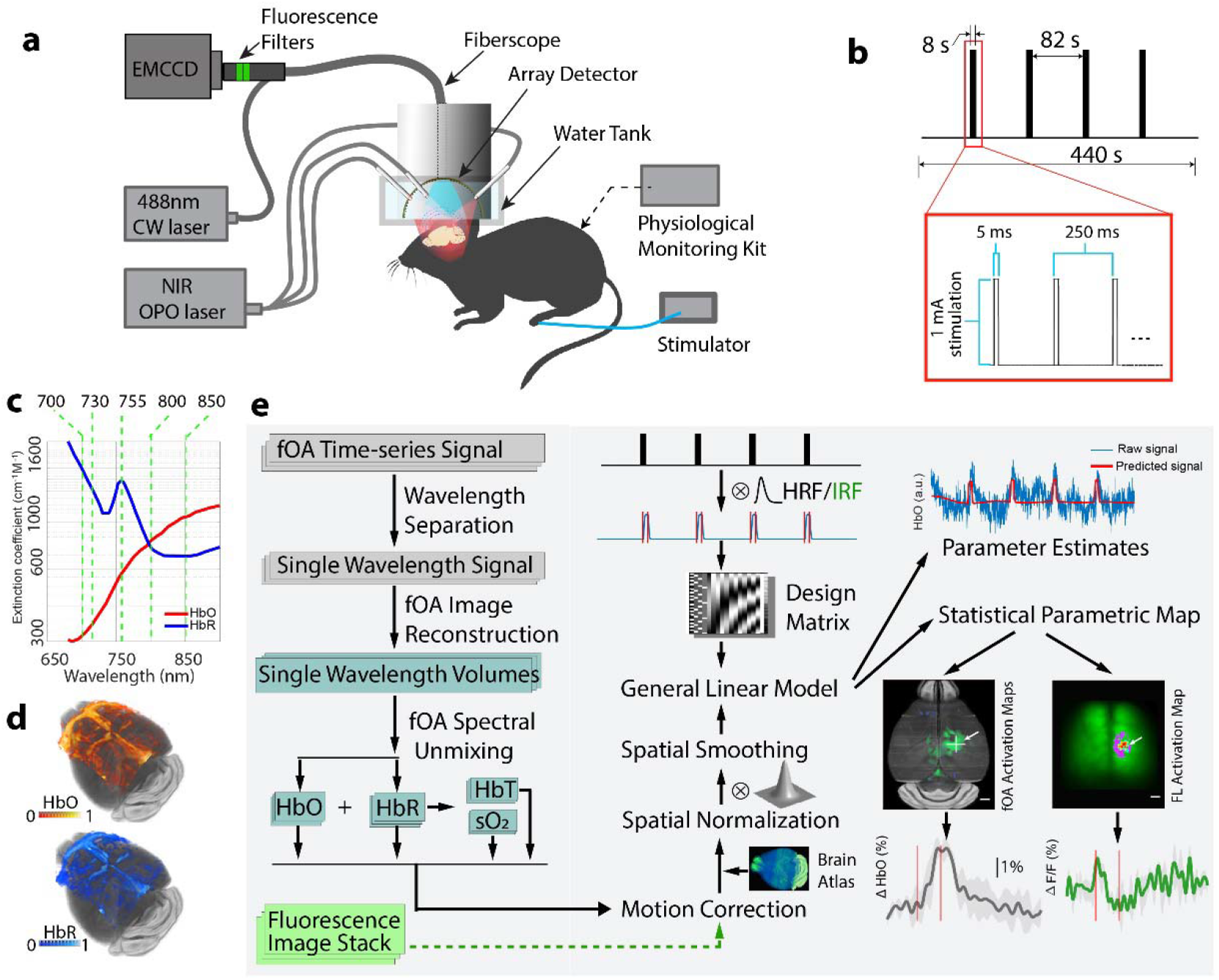
Layout of the hybrid FLOT platform for multi-parametric imaging of murine brain activation. **a** Schematics of the imaging system incorporating the simultaneous functional optoacoustic (fOA) and fluorescence readings. **b** Burst electric current stimulation paradigm applied to the left hindpaw. **c** Wavelength selection for the spectroscopic fOA imaging of blood oxygenation. The molar extinction spectra for HbO and HbR are plotted. **d** Spatial distribution of the spectrally unmixed HbO and HbR hemodynamic components in the brain. **e** The custom FLOT data processing pipeline includes fOA image reconstruction and spectral unmixing (left column), fluorescence data reshaping, pre-processing and the general linear model (middle column), and statistical parametric mapping (right column).

### GCaMP6f impulse Calcium Response Function (CRF)

We first estimated the CRF of the rapid calcium activity using epifluorescence recordings acquired at a high frame rate of 200 Hz in Thy1-GCaMP6f mouse brains *in vivo*. The high frame-rate acquisitions were employed to extract well-defined response function profiles subsequently used in the fluorescence data processing pipeline. The scalp of Thy1-GCaMP6f mice was removed prior to imaging while keeping the skull intact (Fig. 2**a**), which resulted in clearly resolvable cortical vasculature. Temporal fluorescence signal profiles from two representative 0.3 × 0.3 mm^2^ regions of interest (ROIs) were tracked in both the contralateral (opposite side from the stimulated paw) and ipsilateral hemispheres, as indicated in Fig. 2**a**. The recorded profiles were averaged across different stimulation bursts to reveal the activation time course (Fig. 2**b**). The averaged contralateral and ipsilateral signal profiles manifested a high degree of correlation, consisting of both spontaneous wave activity as well as sharp contralateral activity spikes evoked by the electric pulses. The respective averaged time-lapse images further reveal the spatio-temporal patterns of stimulus-evoked neuronal activation in the somatosensory cortex (Fig. 2**c**). In order to estimate the average stimulus-evoked fluorescence response (Fig. 2**d** and Fig. S2), we first applied a temporal notch filter with 0.2 - 3 Hz stop-band to the unaveraged time profiles in the selected contralateral ROI. Next, the filtered profiles were averaged for all the stimulation electric pulses with the resulting curve fitted to a gamma function *y* = *ct*^*a*-1^*e*^−*t*/*b*^ / [*b*^*a*^ *Γ(*a*)], a = 4.775, b = 0.016, c = 0.079. In the subsequent analysis, the gamma fitted curve was adopted as the characteristic GCaMP6f CRF to external electrical paw stimulation. The calculated correlation coefficient between the measured curve and the fitted gamma function is 0.991 with p-value < 10^−5^ for t test. Other functional parameters were further calculated across all the stimulation trials, including the activation intensity 1.3 ± 0.4%, time-to-peak (TTP) 0.063 ± 0.013 s, full-width-at-half-maximum (FWHM) of the calcium response 0.085 ± 0.024 s and decay time (T_1/2_) 0.106 ± 0.025 s (Fig. 2**e**). A linear regressor was then obtained by convolving the stimulation paradigm with the GCaMP6f CRF. Fluorescence data processing with the proposed pipeline rendered an activation map revealing the expected location of the activated region with high specificity (Fig. 2**f**). Compared to an activation map based on the difference image between activated frames and baseline recording, the proposed method exhibits superior robustness and has better localization specificity as it is less affected by the slow changes in light intensity affected by hemoglobin absorption variations (Fig. S3).

**Figure 2.**
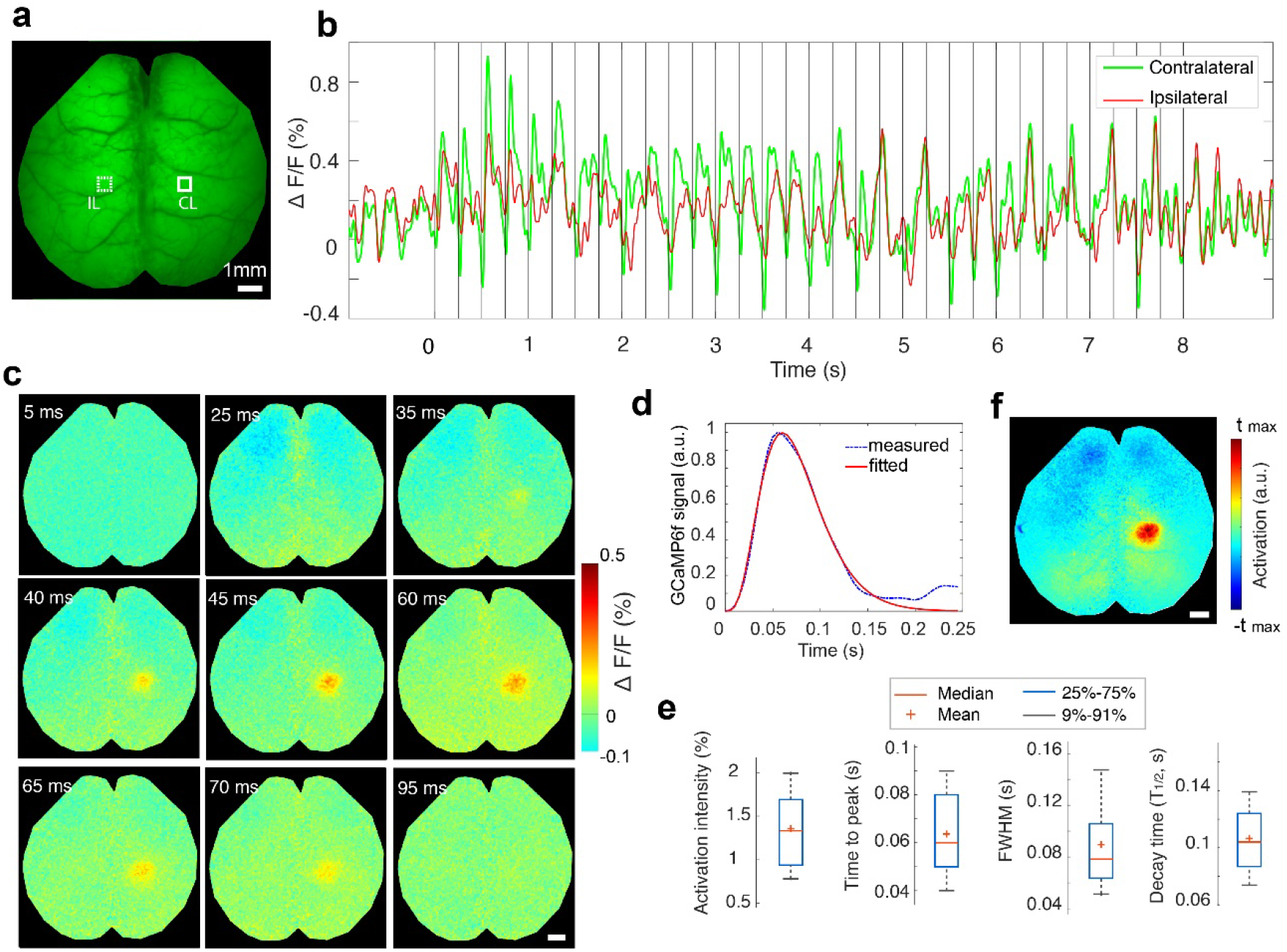
Assessment of the GCaMP6f impulse response function to electrical paw stimulation in mice. **a** Representative time-averaged epi-fluorescence image of the mouse brain. **b** Temporal fluorescence signal profiles spatially averaged over two 0.3 × 0.3 mm^2^ sized ROIs in the contralateral (CL) and ipsilateral (IL) somatosensory cortex, as indicated in panel **a**. Each vertical line indicates single electric pulse. **c** Time-lapse images post the pulsed electric current stimulation obtained by averaging the responses from all the stimulation trials (pulses). **d** Averaged stimulus-evoked response function estimated by averaging activation curves from 160 consecutive stimulation pulses fitted to a Gamma function. **e** Functional parameters extracted from all the stimulation trials. **f** Activation map obtained from the proposed data processing pipeline. All scale bars: 1 mm.

### FLOT reveals concurrent calcium and hemodynamic response maps

Next, we simultaneously imaged calcium and hemodynamic responses in Thy1-GCaMP6f mice (n = 6). Since the fOA imaging was performed in the near-infrared spectral window, the scalp was kept intact for 4 mice while it was removed for another 2 mice to mitigate image artifacts due to skin pigmentation (see Methods section). Activation maps corresponding to different hemodynamic components, namely HbO, HbR, total hemoglobin (HbT) and oxygen saturation (sO_2_), were further rendered from the fOA image sequences. Localized responses were clearly observed in the transverse, sagittal and coronal cross sections overlaid to the Allen P56 mouse brain atlas^13^ (Fig. 3**a**; voxels in the activation map are statistically significant with respect to a null hypothesis of no activation: p < 0.05 one-sample t-test after false discovery rate (FDR) correction). Localized activity was detected within the primary somatosensory cortex as well as within the primary motor area on the contralateral side. No obvious activation was observed in the corresponding regions on the ipsilateral side. Since the HbR response was negatively correlated to the hemodynamic response regressor while other components were positively correlated, both positive and negative thresholds were applied to the calculated activation map (i.e., t-map) of each component. Multiple functional parameters were estimated from these fOA maps, i.e., the activation intensity, TTP, FWHM, decay time and activation volume size (Fig. 3**b**).

**Figure 3.**
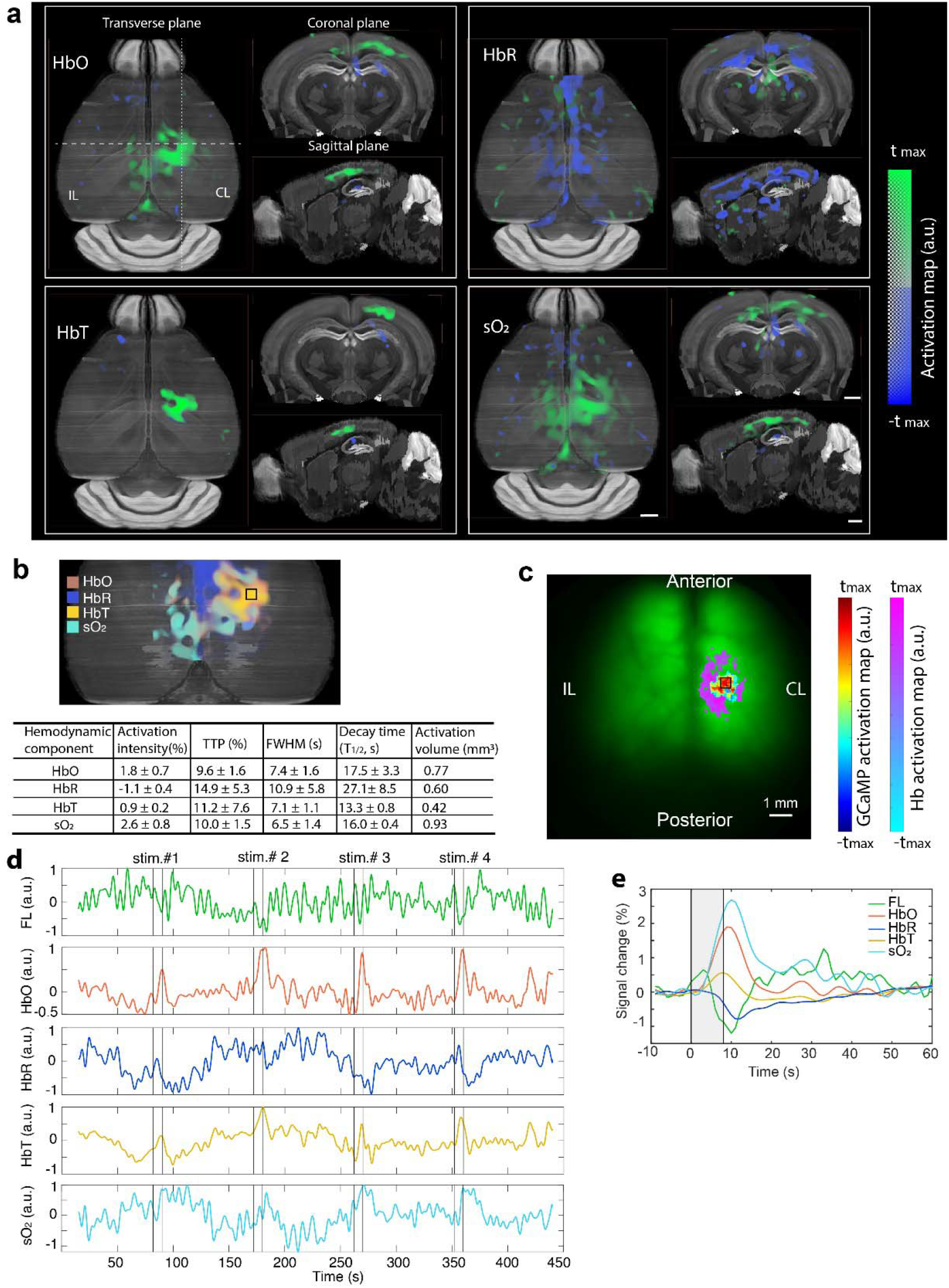
Concurrent measurement of calcium and hemodynamic responses in the mouse brain. **a** Transverse, sagittal and coronal views of the activation maps corresponding to HbO, HbR, HbT and sO_2_ components extracted from fOA images. **b** Corresponding GCaMP and hemodynamic activation maps calculated from the fluorescence measurements overlaid with the fluorescence image of a GCaMP mouse brain with intact skull and scalp. **c** Superimposed fOA activation maps and statistics of the functional parameters from the ROI indicated by the black square situated at a depth of ^~^300 μm from the surface of the cortex. **d** Unaveraged time courses of the fluorescence and fOA signals from the ROIs indicated in **b** and **c**. The fluorescence signal is lowpass filtered to emphasize slow trends (top row; Butterworth, cutoff frequency 0.5 Hz). **e** Averaged activation time courses of fluorescence, HbO, HbR, HbT and sO_2_ from the ROI indicated in **b** and **c**. CL: contralateral, IL: ipsilateral. All scale bars: 1 mm.

We next analyzed the concurrently-acquired fluorescence data. Due to the strong light scattering from the scalp, the fluorescence images appeared blurry compared to those acquired after scalp removal. Yet, localized brain activation upon electrical paw stimulation was clearly observed through the intact scalp. Moreover, increased light absorption by hemoglobin results in modulation of fluorescence signals by the hemodynamic responses in the activated region. A secondary hemodynamic activation map was thus obtained simultaneously from the fluorescence measurement by simply replacing the GCaMP CRF with the HRF in the construction of the regressor and overlaid onto the GCaMP activation map (Fig. 3**c**; both activation maps are thresholded at 70% of their corresponding maximum t-value).

Both unaveraged fluorescence and fOA signal time courses are shown in Fig. 3**d**, depicting the highly dynamic spontaneous fluctuations and stimulus evoked responses. Significant correlation exists between the simultaneously recorded fluorescence and HbO/HbR responses, despite the fact that the signals were measured with two independent modalities, which may partially explain the slow background fluorescence signal fluctuations. These dynamics may further depend on the physiological status of the mouse, which is profoundly affected by many factors such as anesthesia and individual differences. Fractional averaged signal changes from representative ROIs in contralateral hemisphere are shown in Fig. 3**e**. Interestingly, the lowpass filtered fluorescence signal peaked right after the paw stimulation due to the fast calcium-mediated response but then slowly declined below its baseline value gradually going back to the baseline, reflecting changes in light attenuation associated to increased local hemoglobin absorption in the activated regions.

Our results show that fOA provides diverse information on brain activation responses complementing the fluorescence readings. Further, it was observed that fractional hemodynamic signal changes significantly differ across various brain regions whereas fluorescence responses did not exhibit such spatial variability (Fig.S4).

### Multi-parametric analysis of stimulus-evoked brain activation

Multi-parametric analysis was performed to characterize the coupling between simultaneous fluorescence and fOA readings. The same data processing pipeline was repeated for all mice (n = 6) across a total of 80 stimulation trials. Activation maps were first calculated using both the fluorescence and fOA images to infer activated brain regions and thereby to select ROIs for calculating the time courses. Spontaneous neural activity largely averages-out pre- and post-stimulation while the stimulation evoked responses are preserved (Fig. 4**a**). The fractional signal changes of the different hemodynamic components were then averaged across animals (Fig. 4**b-e**, n = 6). Strongly correlated signal increase in HbO (Fig. 4**b**), HbT (Fig. 4**d**) and sO_2_ (Fig. 4**e**) and HbR decline (Fig. 4**c**) were observed from the fOA measurements. As expected, these signal changes are closely correlated to the predicted hemodynamic response by convolving the HRF with the mean neural activity trace (green curve in Fig. 4**a**). Based on the fractional change in HbO and HbR, the fluorescence signal contaminated by slowly varying changes in hemoglobin absorption was successfully corrected^5^ using estimates of the pathlength factors xEx = 0.56 mm^−1^ and xEm = 0.57 mm^−1^ (Fig. 4**f**).

**Figure 4.**
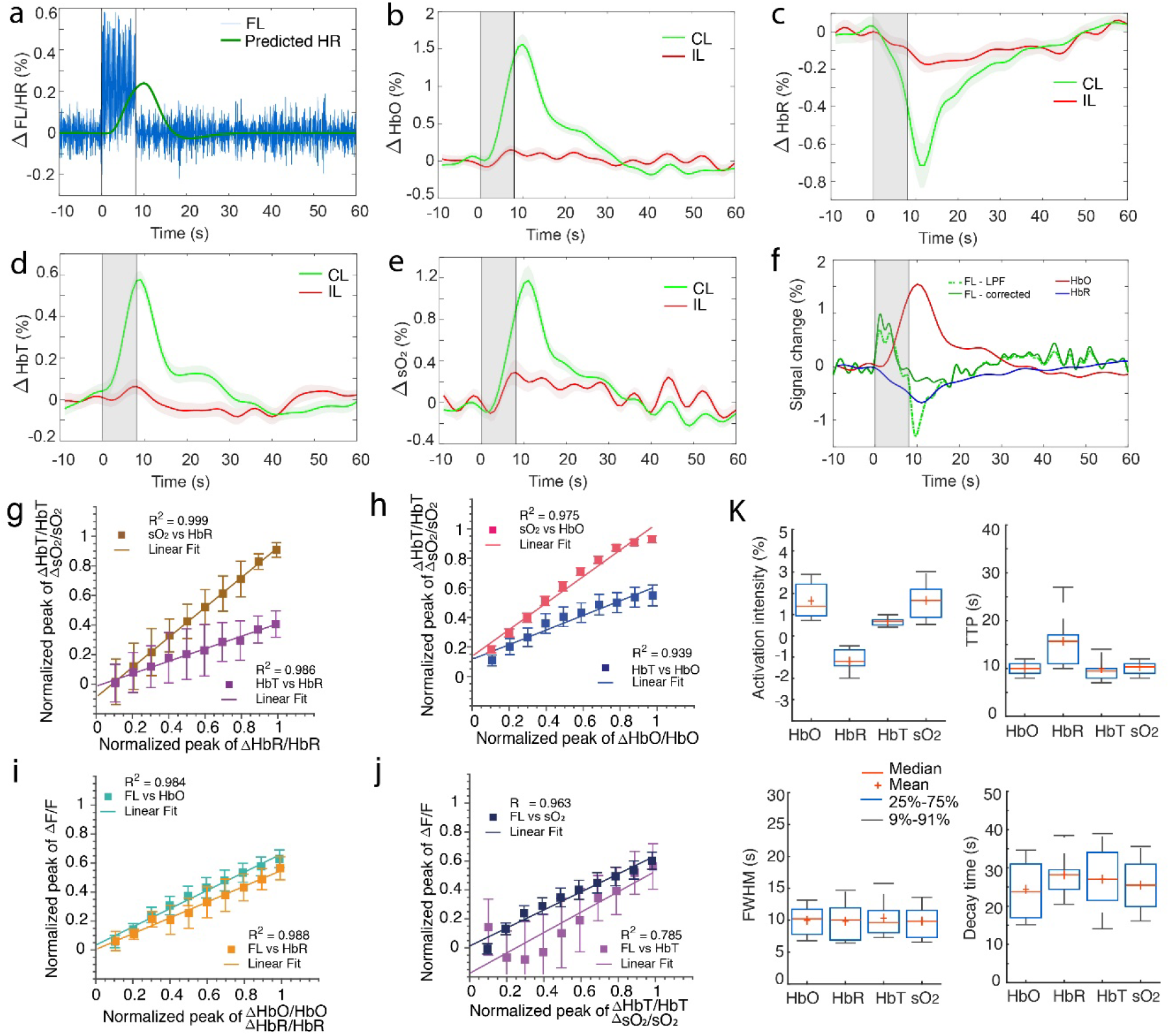
Multi-parametric analysis of the coupling between simultaneous fluorescence and fOA readings. **a** Averaged fluorescence activation curve after applying a bandpass filter (3-8 Hz) and the predicted hemodynamic response (HR) calculated via convolution with the HRF. **b-e** Averaged activation curves of HbO, HbR, HbT and sO_2_ across all the stimulation trials. Shaded regions show standard error of mean (SEM) across n = 6 mice in all the trials. **f** Fluorescence signal compensated for hemoglobin absorption variations by considering the fractional changes of HbO and HbR from the contralateral side. **g-j** Coupling between HbO, HbR, HbT, sO_2_ and lowpass filtered fluorescence signal from simultaneously measured single activation traces. **k** Statistics of the activation intensity, TTP, FWHM and decay time (T_1/2_) across all the trials from 6 mice. Note that all values are statistically significant compared to their ipsilateral counterparts (p-value < 0.05, two-tailed paired t-test). CL: contralateral, IL: ipsilateral.

The coupling between neural activity and the various hemodynamic response components was further studied by calculating the relationship between the peak values of the corresponding (simultaneously measured) single activation traces of different normalized response components (Figs. 4**g-j**). In Figs. 4g-h, the results show a strong coupling relationship between HbT and sO_2_ responses vs. both HbR and HbO responses, respectively. These results show that both HbT and sO_2_ responses saturate for high HbO responses, while they’re uniformly linear across the full HbR response range. Further, by comparing the ranges and standard deviations between the different plots it is seen that both HbT and sO_2_ are more closely coupled with HbO than with HbR. Besides, an overall stronger coupling of sO_2_ responses to both HbO and HbR was observed. The strong relationship between the different hemodynamic components extracted from the fOA measurement and the fluorescence signal corroborates the synergistic detection of neurovascular coupling phenomena in stimulus-evoked brain activation by the hybrid FLOT method.

Despite the strong correlation among fOA components, noticeable differences were observed in the calculated functional parameters (Fig. 4**k**). sO_2_ (1.7 ± 0.9%) presents stronger activation intensity as compared to HbO (1.6 ± 0.9%), HbR (−1.2 ± 0.6%) and HbT (0.7 ± 0.2%). In terms of TTP, the HbR readings (15.7 ± 5.9 s) exhibit the most delayed response followed by sO_2_ (10.3± 1.3 s), HbO (10.3 ± 1.4 s) and HbT (9.9 ± 2.9 s). Similar pattern was observed in the FWHM values with HbT (11.5 ± 4.9 s) holding the longest response duration followed by HbR (11.3 ± 6.5 s), HbO (10.5 ± 4.0 s) and sO_2_ (10.3 ± 3.8 s). In terms of decay time (T_1/2_), the HbR (28.2 ± 5.9 s) holds the longest value followed by HbT (27.0 ± 8.6 s), sO_2_ (25.5 ± 6.7 s) and HbO (24.4 ± 7.3 s).

## Discussion

Major efforts are underway to devise new neuroimaging methods enabling scalable recording of neural activity and the associated hemodynamic responses. Both calcium and hemodynamic changes can be captured with sequential (non-concurrent) multi-modal strategies combining different contrast mechanisms. However, the mammalian brain’s complexity and its highly dynamic physiological status challenge the establishment of interpretable links across separately acquired multi-modal readings often covering different spatio-temporal scales. In this work, calcium signal and hemodynamic responses were imaged concurrently and a tailored functional data processing pipeline was developed to facilitate the comparison of calcium and hemodynamic responses. Significant correlation was revealed between the GCaMP fluorescence signal and the associated increases in the HbO, HbT, sO_2_ and decrease in the HbR signals. In previous neuroimaging studies with fMRI, it was determined that HbR changes are the chief contributors to the BOLD signal. Interestingly, this signal is affected by changes in blood flow, blood volume, and metabolic rate of oxygen, which makes the BOLD signal relatively ambiguous to interpret. In contrast, we were able here to retrieve signal changes from individual hemodynamic components simultaneously. The cross-correlation analysis (Figs. 4**g, h**) indicates that both ΔHbO and ΔHbR have comparable contribution to the local change in HbT and sO_2_ while ΔHbO has slightly higher contribution to ΔHbT. The coupling between calcium signal and hemodynamics is further corroborated by cross-correlation between ΔHbO/ΔHbR and the hemodynamic changes ΔHb predicted by analyzing the GCaMP signal convolved with the HRF. These findings complement previous observations on event-evoked brain activation mapping and related hemodynamic readings based on wide-field optical mapping and fMRI^2,5,14–17^. The hybrid FLOT approach expands the capabilities of previously reported multi-modal methods to provide more comprehensive functional information on stimulus-evoked neural activation in the murine brain.

fOA is particularly advantageous for characterizing hemodynamic responses in the brain as it can map multiple hemodynamic parameters in 3D with high spatial resolution and ultrafast (real-time) volume rate. Recently it has been shown to be sensitive to fast stimulus-evoked calcium responses mediated by spectral absorption changes in GCaMP proteins^18,19^. However, accurate spectral unmixing of GCaMP-associated changes from the strong background absorption by hemoglobin remains challenging with fOA, especially when employing calcium-sensitive molecules with peak absorption in the visible spectral range. We have considered the possible confound where spectral unmixing of HbO and HbR with a linear model leads to quantitative errors in the sO_2_ and HbT readings due to the wavelength-dependent light attenuation in the tissue^18,19^. It has previously been shown that accurate estimation of oxygenation can be obtained by selecting a set of optical wavelengths that (1) minimize spectral coloring effects, (2) avoid ill-conditioning in the inverse problem, and (3) provide a sufficiently high SNR^21^ The wavelength combination in this work was carefully chosen to ensure optimal spectral unmixing of the fOA images. Specifically, we avoided the visible spectral range of 500 – 600 nm, where significant spectral coloring associated to the strongly wavelength-dependent blood absorption is produced. Furthermore, one of the excitation wavelengths was selected at the isosbestic point of hemoglobin (800 nm), another wavelength at the fingerprint spectral point of HbR (755 nm) and the remaining three wavelengths of 700, 730 and 850 nm exhibiting large absorption difference between HbR and HbO. This ensures a small condition number in the molar absorption coefficient matrix. NIR wavelengths are known to more efficiently penetrate biological tissues, thus providing a good balance between penetration depth and SNR for deep brain imaging. Note however, that fOA generally provides inferior penetration depth as compared to MRI or positron emission tomography, while the high resolution of the method can be further compromised by the skull-induced ultrasound distortions^22^.

Accurate estimation of the HRF shape is essential to preventing both false positives and false negatives that may result from a mismatch between the predicted and measured signals. Considering that no specific HRF model currently exists for the different hemodynamic readings provided by fOA, we employed a modified SPM canonical HRF with a TTP of 2.3 s and FWHM of 1.9 s to represent the rodent hemodynamic response^23,24^. This simplified model was shown to effectively infer active voxels in the mouse brain. Note, however, that the shape of the HRF has been shown to be affected by the particular selection of tasks and brain regions, thus limiting its general applicability. Also, the hemodynamic response to a brief stimulus in the rodent brain is species-dependent and can significantly change during development^17^. Ideally, the HRF parameters should be directly interpretable in terms of changes in neuronal activity and should be estimated so that statistical power is maximized. However, substantial differences among models in terms of power, bias and parameter mismatches have been reported in fMRI studies^25^, which are similarly expected to affect the fOA measurements. Hence, more sophisticated HRF models optimally fitting the different hemodynamic components are expected to bring more flexibility in estimating the functional parameters, which will be addressed in future work.

In summary, the unique capabilities of FLOT for simultaneous imaging of fast calcium signals associated to neural activity and the induced hemodynamic responses across the entire mouse cortex noninvasively can reveal new insights into the basic mechanisms and extent of stimulus-evoked responses. fOA is a versatile tool to resolve multiple hemodynamic parameters in real-time, thus providing additional information that may enhance or complement existing methods of studying the neurovascular and neurometabolic coupling mechanisms. Epifluorescence has high sensitivity and spatio-temporal resolution when it comes to tracking calcium activity in the cortical regions, which served to corroborate and complement the fOA observations. The fOA readings were in turn used to decouple the calcium responses from the slowly varying hemodynamic variations in the fluorescence recordings. The proposed approach does not require cranial windowing or any other invasive surgical procedures. It can also be used in a fully non-invasive manner in mice with non-pigmented scalps. All in all, FLOT is a powerful new brain imaging tool that can be broadly applied for investigations into neurovascular coupling, cerebrovascular and neurodegenerative conditions as well as monitoring of therapies.

## Methods

### Hybrid epifluorescence and optoacoustic tomography (FLOT) system

Hybrid epifluorescence-optoacoustic imaging was achieved with a system that integrates a spherical matrix array transducer (Imasonic SaS, Voray, France) for volumetric fOA imaging with a custom-made fiberscope (Zibra Corporation, Westport, USA) for co-registered optical (epifluorescence) imaging (Fig. 1**a**). The transducer array consists of 512 piezocomposite elements arranged on a hemispherical surface with a 150° angular coverage (1.48π solid angle). Individual elements have 7 MHz central frequency and >80% detection bandwidth. The array features a central 8 mm diameter aperture and three additional 4 mm diameter apertures located at 45° elevation angle and equally spaced in the azimuthal direction. The fiberscope consists of a 1.4 mm diameter optic image guide consisting of 100,000 fibers collecting the fluorescent responses and an illumination bundle composed of 19 fibers having 600 μm core diameter and 0.4 numerical aperture (NA) for optical excitation (Zibra Corporation, Westport, USA). Epifluorescence imaging was performed by inserting the tip of the fiberscope into the 8 mm aperture of the matrix array. At the output end of the image guide, two emission filters (FL514.5-10, Thorlabs, USA) were cascaded to isolate the fluorescence emission from GCaMP, which was subsequently captured with an EMCCD camera (iXon Life, Andor, UK) at 20 fps. The hybrid FLOT system provided a co-registered 12 mm diameter FOV with 40 μm lateral resolution for fluorescence and nearly isotropic 3D spatial resolution down to 113 μm for fOA imaging. Note that for characterizing the GCaMP6f impulse response with high temporal resolution in the millisecond range, the EMCCD camera having superior SNR was replaced with a high-speed CMOS camera (PCO.dimax S1, PCO, Germany) operating at 200 fps. Fluorescence excitation was provided with a continuous wave 488 nm laser (Sapphire LPX 488-500, Coherent, USA). On the other hand, a custom-made fiber bundle (CeramOptec GmbH, Germany) was used to guide a short-pulsed (<10 ns) beam generated with an optical parametric oscillator (OPO) laser (Spit-Light, Innolas Laser GmbH, Germany) through the three lateral apertures of the array. The laser wavelength was rapidly swept between five wavelengths (700 nm, 730 nm, 755 nm, 800 nm, and 850 nm) on a per-pulse basis at 100 Hz pulse repetition frequency (PRF). The pulse energy was ^~^11 mJ at the output of the illumination fiber bundle. The generated signals were acquired with a custom-made data acquisition system (DAQ, Falkenstein Mikrosysteme GmbH, Germany) at 40 Megasamples per second (Msps) and recorded raw data transmitted to a PC via Ethernet. Synchronization of the excitation light pulses, the fluorescence and fOA data acquisition, and the electrical paw stimulation was achieved with an external trigger device (Pulse Pal V2, Sanworks, USA).

### Animal models

Female GCaMP6f mice (C57BL/6J-Tg(Thy1-GCaMP6f) GP5.17Dkim/J, the Jackson Laboratory, USA, 6 to 11 week-old, n = 6) were employed in this study. Animals were housed in individually ventilated, temperature-controlled cages under a 12-hour dark/light cycle. Pelleted food (3437PXL15, CARGILL) and water were provided ad-libitum. All experiments were performed in accordance with the Swiss Federal Act on Animal Protection and were approved by the Cantonal Veterinary Office Zurich.

### *In vivo* imaging

Mice were anesthetized for the *in vivo* imaging experiments. Anesthesia was inducted with intraperitoneal injection of ketamine (100 mg/kg body weight, Pfizer) and xylazine (10 mg/kg body weight, Bayer) cocktail. The bolus injections were separated into two injections with 5 min gap to prevent cardiovascular complications. Maintenance injection was administrated every 45 min with a mixture of ketamine (100 mg/kg body weight) and xylazine (2.5 mg/kg body weight). Both the scalp and skull of the mice were kept intact (n = 4) on the premise that no pigment in the scalp were presented while the scalp was removed and skull was kept intact for mice with scalp pigmentation (n = 2). Scalp removal was performed after injection of Buprenorphine (0.1 mg/kg body weight, Temegesic, Indivior, Switzerland) together with hemostatic sponges (Gelfoam®, Pfizer Pharmaceutical) to minimize bleeding. Imaging was performed by placing each mouse onto a 3D translation stage in a prone position that facilitated optimal positioning of the region of interest in the brain in the center of the FOV of both modalities. Ultrasound gel was applied on the mouse head to ensure optimal acoustic signal coupling. The mouse head was immobilized using a custom stereotactic frame (Narishige International Limited, London, United Kingdom). During the experiment, an oxygen/air mixture (0.1/0.4 L.min^−1^) was provided through a breathing mask. Peripheral blood oxygen saturation, heart rate and body temperature were continuously monitored (PhysioSuite, Kent Scientific) during data acquisition. The body temperature was kept around 37 °C with a feedback-controlled heating pad.

A BOLD fMRI comparable stimulation paradigm in terms of stimulation duration and intensity was adopted in this study^15^. Unipolar rectangular electric pulses of 5 ms duration and 1.0 mA intensity were applied to the left hindpaw at 4 Hz stimulus frequency, 8 s onset time, and 82 s burst intervals (Fig. 1**b**). The electric signals were generated using a stimulus isolator device (Model A365R, World Precision Instruments, USA) fed by an external trigger (Pulse Pal V2, Sanworks, USA). For each sequence, the total duration of the stimulation pulse train was 440 s with the individual stimuli synchronized with the excitation light and data acquisition system. The sequence was repeated 2-5 times per animal. After the experiments, the animals with scalp removed were euthanized while still under anesthesia, whereas other mice with intact scalp emerged from the anesthesia and fully recovered.

### Custom data analysis pipeline for both fluorescence and fOA

A data analysis pipeline inspired by SPM12^26,27^ was developed for both fluorescence and fOA data analysis (Fig. 1**e**). To this end, a custom script was developed with MATLAB (2019b, MathWorks, USA). A general linear model (GLM) was applied to a voxel-by-voxel based statistical analysis. fOA volumetric data and fluorescence images were pre-processed before being fed into the model. Specifically, fOA raw signal matrices were first separated according to the corresponding excitation wavelengths and the signals bandpass filtered between 0.1-8 MHz. The images were reconstructed separately for each wavelength using a filtered back-projection algorithm^28^ (100×100×100 μm^3^ voxel size, 8×8×4 mm^3^ FOV). Generally, the fOA signal at an arbitrary point ***r*** within the imaged tissue volume is mainly resulting from the light absorption by HbO and HbR, which can be approximated as^29^

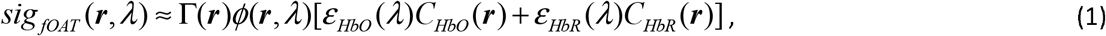

where Γ is the Grueneisen parameter, λ is the wavelength of the illumination laser beam, ***ϕ*** is the light fluent within the tissue and ***ε*** and *C* are the molar extinction coefficients and concentrations of each absorbing substance (molecule), respectively. fOA signals were firstly normalized to the laser pulse energy at each wavelength. To reduce the spectral coloring effects^21^, i.e., the spatially and wavelength dependent light fluence distribution within the tissue, an exponential light attenuation model was applied to compensate for the depth-dependent signal decay. The effective attenuation coefficient (μ_eff_) was calculated as^30^

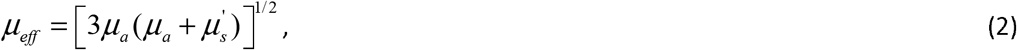

where *μ*_a_ is the absorption coefficient and was set to 0.1, assuming homogenous light absorption in the mouse brain at all the 5 wavelengths. 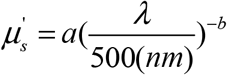 is the reduced scattering coefficient whereas a=24.2 cm^−1^ and b= 1.611 were assumed based on the averaged scattering properties of the brain tissue ^30^.

After light fluence correction, the concentrations of HbO and HbR for a spatially independent Γ were estimated via linear spectral unmixing as

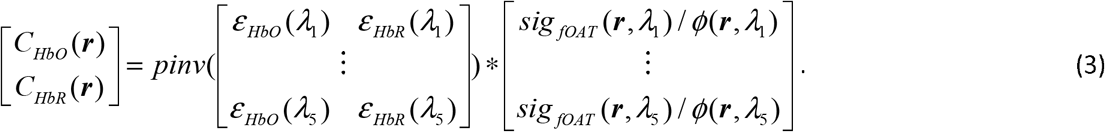

Subsequently, HbT and sO_2_ were calculated from the concentrations of HbO and HbR as

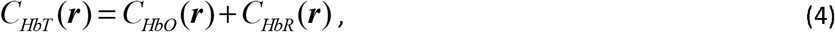

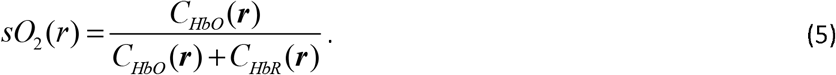

For fluorescence image pre-processing, the image stack was rearranged to form a 4D volume with the third dimension equal to 1 to make it compatible to the overall processing pipeline. Motion estimation and correction was performed by using SPM12 realign module before the processed data was fed to the model for statistical analysis. The custom stereotactic frame helped mitigating the motion artifacts. The relative center of mass displacement in each trial typically remained below 0.2 voxel, thus ensuring negligible motion-induced errors (false activations). Spatial smoothing (Gaussian kernel FWHM= 0.3 mm) was further applied to reduce random fluctuations and reconstruction-related image artifacts. A high-pass filter of 1/135 Hz was then used to detrend the fOA data. For fluorescence data, a lowpass filter with 0.5 Hz cutoff frequency was applied to retrieve the slow hemodynamic response encoded in the fluorescence signal and a bandpass filter (3 - 8Hz) was applied to extract the fast calcium transients.

In the GLM, the regressor was obtained by convolving the stimulation paradigm (Fig. 1**b**) with either the measured GCaMP CRF (for fluorescence data) or a modified SPM canonical HRF (for fOA data) having a TTP of 2.3 s and FWHM of 1.9 s to represent the rodent hemodynamic response^23,24^. The regressor together with a constant vector formed the design matrix. The GLM is expressed in the matrix format via^31^

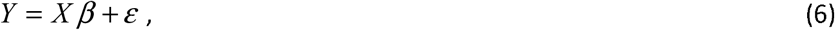

where Y is the column vector of observations representing a signal voxel sampled at successive time point, ε is the column vector of error terms, β is the column vector of parameters and X is the design matrix. The estimated parametric map is calculated as

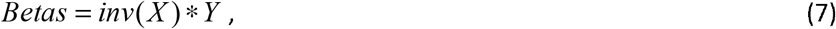

The statistical significance of the event-evoked responses in the observations was further evaluated with a contrast vector c = [1, 0]^T^. The t statistic map (i.e. t-map) and probability levels (i.e. p-values) were calculated as the statistical inferences. Note that for rendering the activation map for fOA, the false discovery rate (FDR) controlling^32^ with p <0.05 was introduced to the t-map. For the fluorescence recordings, the parameteric map (*Betas* in Eq. (7)) was used directly to infer the activation.

The activation map and the corresponding structural image acquired at 850 nm excitation wavelength were overlaid to the Allen Mouse Brain Atlas 2015^33^. The atlas volume has 320 × 456 pixels in each coronal plane and consists of 528 slices with 25 μm isotropic voxel size. Due to the lack of common features and utterly different intensity distribution, unbiased automated registration of fOA with MRI remains challenging^34^. In this work, the commonly used manual landmark based coregistration method^35^ was employed for the alignment in Amira (version 5.4, Thermo Fisher Scientific, USA). Firstly, the structural volumetric data of the mouse brain was reconstructed with the averaged signal matrix acquired at 850 nm. Secondly, this structural volume and the mouse brain atlas were loaded into Amira for pre-alignment. A coarse transformation with respect to translation and rotation was tuned according to the landmarks in both datasets. Finally, the position of the structural optoacoustic volumetric data was finetuned based on cross-sections along the three planes (perspectives) so that landmarks such as the brain curvature and vessel structures matched the MRI brain atlas. The calculated transform information in the transform editor in Amira facilitated registering all the other reconstructed parameters such as HbO, HbR, HbT and sO_2_ from the same mouse to the brain atlas. The overlaid activation map was subsequently rendered and displayed along different planes.

The time courses of brain activation were retrieved from the raw 4D data, namely, 3D volume time series for fOA and 2D image series for fluorescence data. The calculated activation map served to infer responsive ROIs in both contralateral and ipsilateral hemispheres. A time window including 10 s pre-stimulation, 8 s onset and 50 s post-stimulation was selected. Baseline signals of each stimulation cycle were calculated by averaging the 10 s pre-stimulation time window. This was used for calculating fractional signal changes, which were averaged across different stimulation cycles. All the trials acquired at slightly different anesthesia depth and from different mice were averaged to reveal the stimulus-evoked brain activation in a more robust way. Statistics on activation intensity, TTP, FWHM as well as decay time (T_1/2_) were further performed across all the trials. The statistical significance between the averaged values from the contralateral side and their ipsilateral counterparts were evaluated with two-tailed paired t-test.

## Acknowledgements

The authors acknowledge grant support from US National Institutes of Health (UF1 NS107680) and the European Research Council (ERC-2015-CoG-682379).

## Ethics statement

All procedures involving mice conformed to the national guidelines of the Swiss Federal Act on animal protection and were approved by the Cantonal Veterinary Office Zurich.

## Author contributions

ZC and DR conceived the concept and devised the study. ZC and QZ performed the animal experiments. ZC, IG performed image reconstructions and data analysis. XLDB, RN and MR helped with the experiments and provided helpful discussions. SS and DR supervised the work. All authors contributed to writing and revising the manuscript.

