## supplementary file for "Multi-parametric characterization of brain-wide hemodynamic and calcium responses to sensory stimulation in mice"

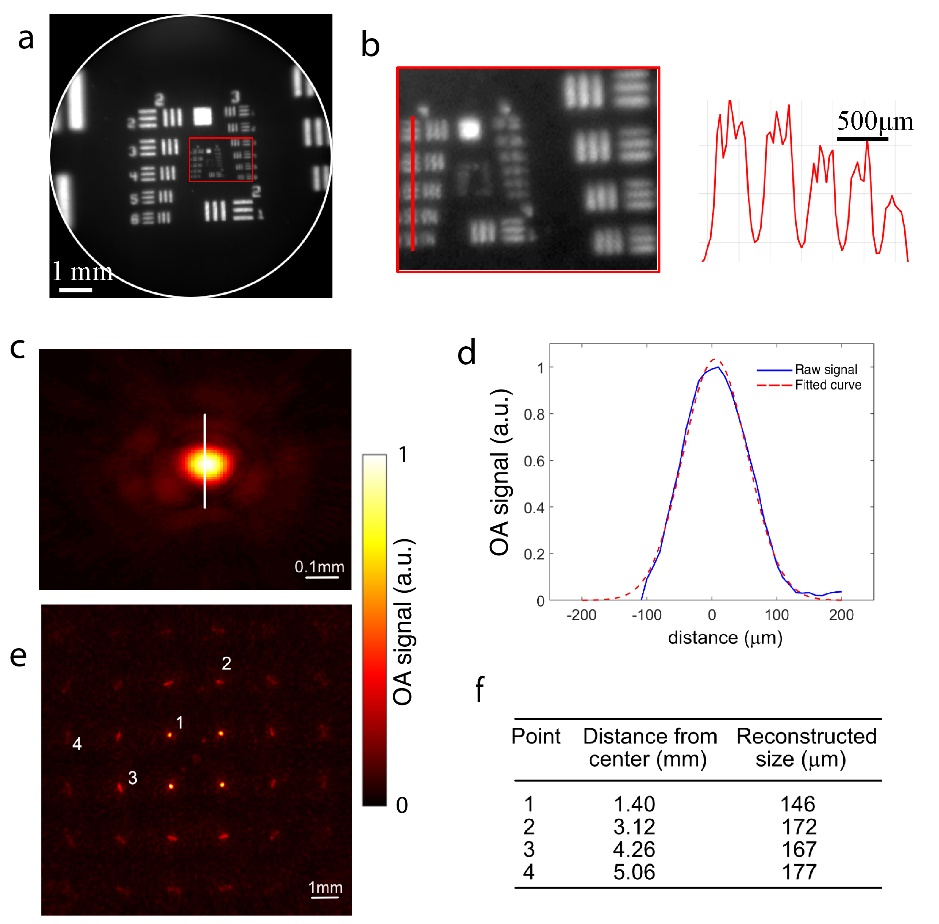

**Fig. S1** Spatial resolution characterization of the hybrid fluorescence optoacoustic tomography (FLOT) system. **a** Image of the 1951 USAF target acquired with a 525 nm bandpass filter. The white circle shows the effective FOV of the fiberscope. **b** Zoomed-in image of the boxed region in **a** and the corresponding 1D profiles along the red lines showing a resolution of 22.62 lp/mm (group 4, element 4). **c** Maximum intensity projection (MIP) along the axial direction of the optoacoustic image of a 30 µm microsphere positioned in the center of the array detector. **d** Vertical profile of the optoacoustic image along the line indicated in **a**. The spatial resolution was estimated via the mean square difference between the width of the fitted curve and the actual microsphere diameter, resulting in a value of 113 µm. **e** A compounded OA image obtained by raster scanning the microsphere across the field of view. **f** The reconstructed microsphere size for positions corresponding to various distances from the center of the spherical array geometry, as indicated in **e**.

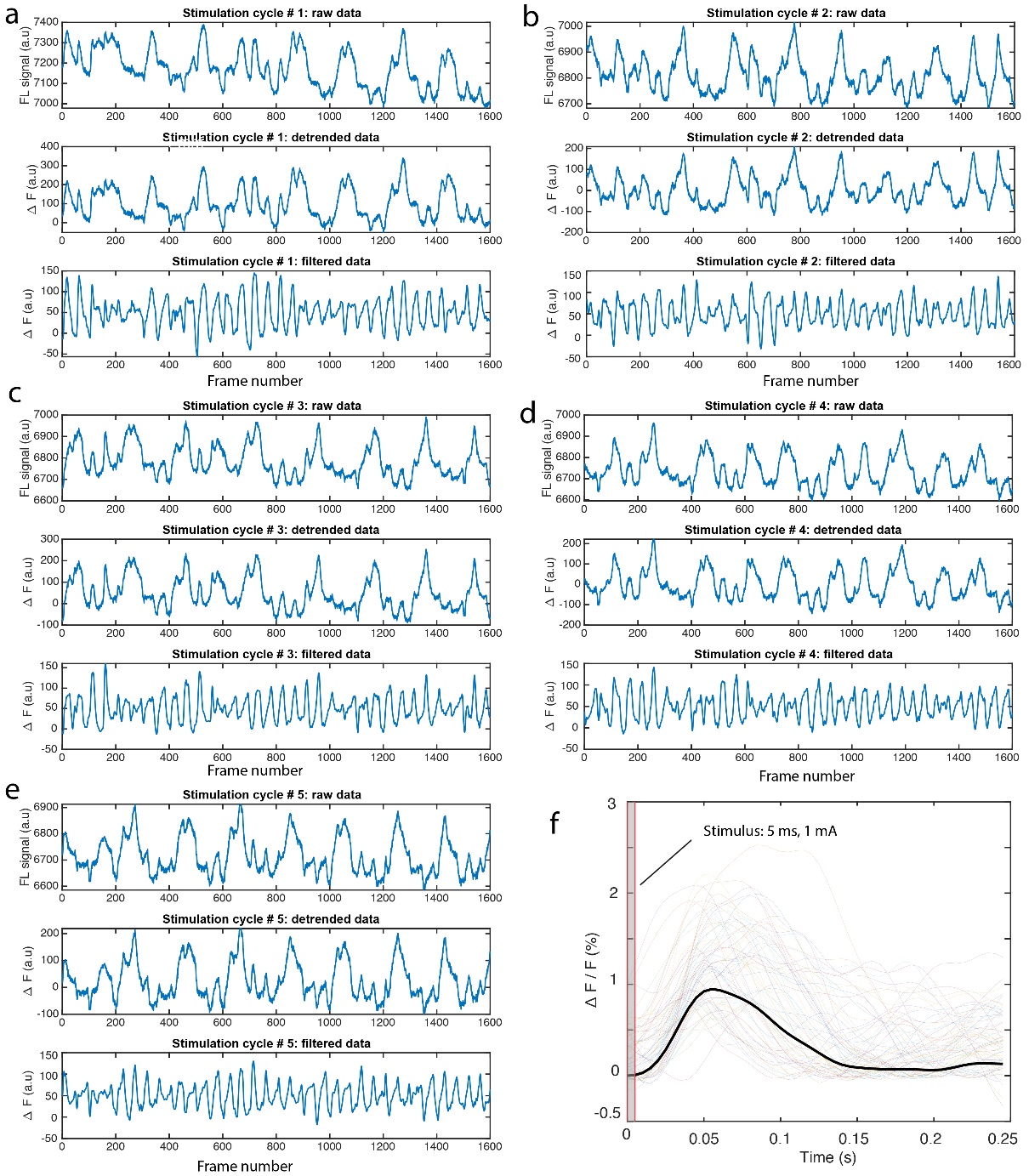

**Fig. S2** Single traces and averaged response from *in vivo* GCaMP6f impulse response measurement. **a** – **e** Signal traces from each stimulation cycle. Top panel: raw fluorescence signal; middle panel: detrended signal; bottom panel: bandpass filtered fluorescence signal with passband of 3-30 Hz. **f** Averaged fractional response overlaid to the 32x5 = 160 traces.

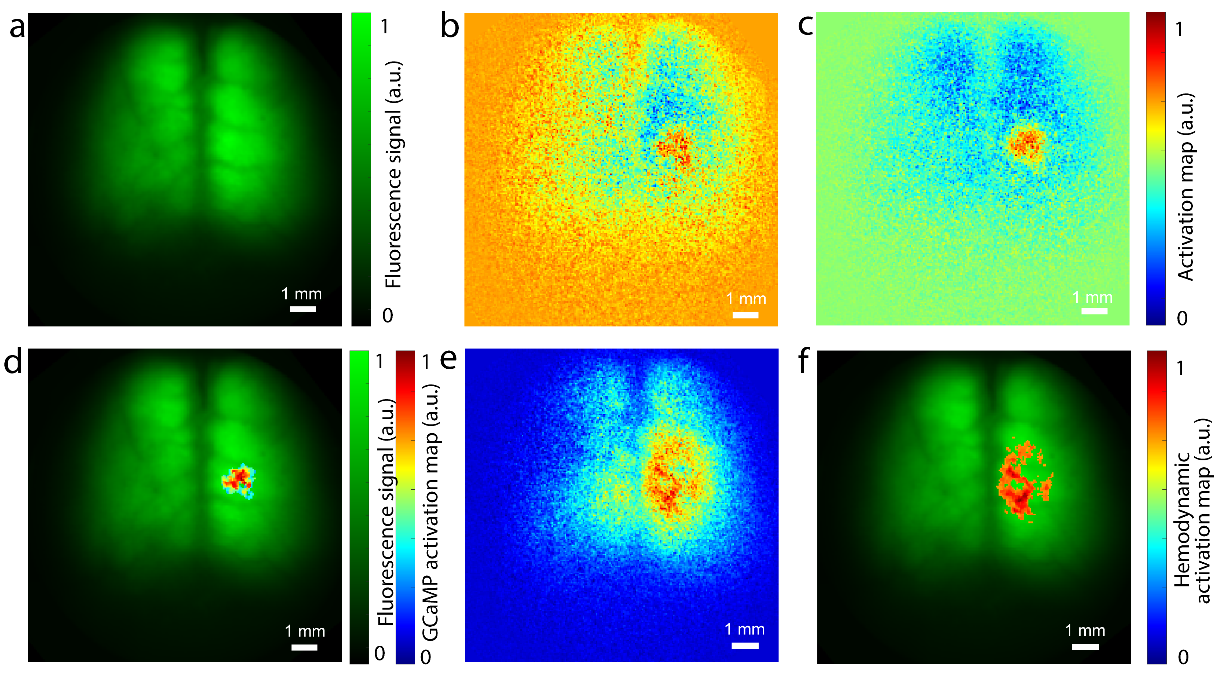

**Fig. S3** Fluorescence experimental results of the GCaMP mouse shown in Fig. 3. **a** Macroscopic fluorescence image. **b** GCaMP activation map calculated by the difference image between the frames acquired during stimulation and before stimulation. **c** GCaMP activation map calculated according to the proposed data analysis pipeline in this work. **d** GCaMP activation map overlaid on the fluorescence image with a threshold of 70% maximum intensity to remove the background. **e** Hemodynamic activation map calculated with the proposed data analysis pipeline in which the GCaMP impulse response function was replaced with the modified hemodynamic response function. **f** Hemodynamic activation map overlaid on the fluorescence image with a threshold of 70% maximum intensity to remove the background.

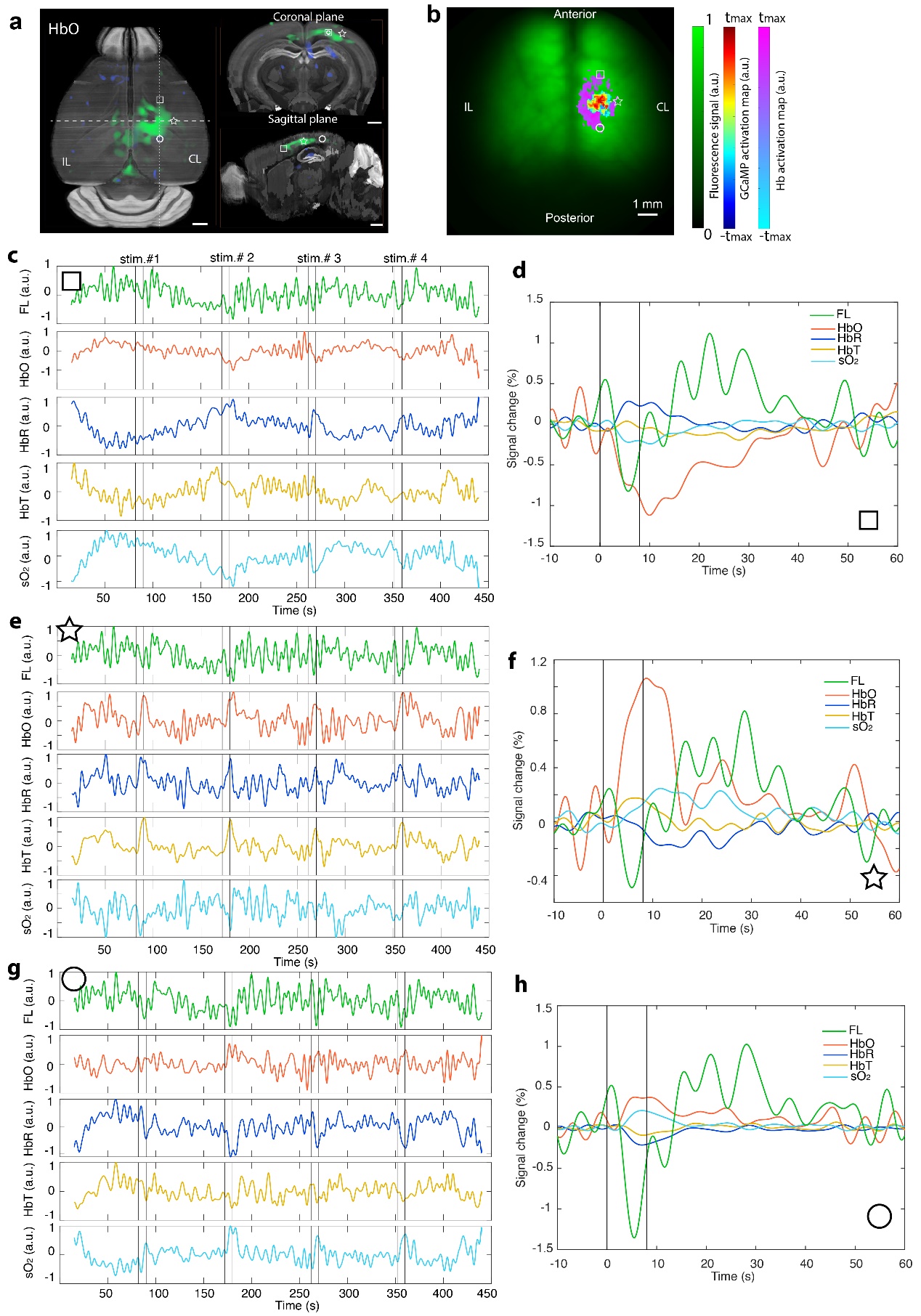

**Fig. S4** Concurrent measurement of calcium and hemodynamic responses in the mouse brain. **a** Transverse, sagittal and coronal views of the activation map from HbO. Regions of interest (ROIs) are indicated by the square, star, and circle for signal analysis. **b** Corresponding GCaMP and hemodynamic activation map along with the ROIs. **c, e, g** Unaveraged time courses of the fluorescence and fOA signals from the ROIs indicated in **a** and **b**. **d, f, h** Fractional signal changes from each ROI by averaging all the four stimulation cycles as shown in **c**, **e,** and **g**, respectively. It is noted that fOA signal changes significantly differ across the listed three brain regions whereas fluorescence responses do not exhibit such spatial variability. CL: contralateral, IL: ipsilateral. All scale bars: 1 mm.
